# Insertions of *P{lacW}* and *P{EP}* artificial transposons on the chromosomal divisions of *Drosophila melanogaster* are not randomly distributed

**DOI:** 10.1101/770172

**Authors:** Bologa Alexandru Marian, Ghita Iulian Cristian, Ratiu Attila Cristian, Ecovoiu Al. Alexandru

## Abstract

Herein we describe the distribution of P{lacW} and P{EP} artificial transposons in the genome of Drosophila melanogaster. A total number of 5,560 P{lacW} and 3,786 P{EP} insertions available on FlyBase were extracted from this database and ordered according to the chromosomal regions they hit. Comparative bioinformatics analysis revealed that the insertion’s patterns are similar for P{lacW} and P{EP}. The two patterns are significantly correlated for chromosomes X, 2L, 2R and 3L, but not for 3R.

Fourier analysis revealed a periodic behaviour in the distribution of insertions, which approximates insertional hotspots. Our results raise questions concerning if and how the local chromosomal landscape affects the insertion patterns of different but related transposons.

## Introduction

Understanding of the insertion sites preferences exhibited by natural transposable elements (TEs) is critical for understanding their role in shaping the host genome structure and evolution. Features of the host genome may contribute to the variation of TEs distributions, such as heterochromatic regions, which are enriched in TEs compared with euchromatic regions. However, the asymmetrical distribution of TEs in the eukaryotic genomes is not well understood (1). In *Drosophila melanogaster*, natural P elements (a class II transposon) are remarkable abundant within the proximal promoter regions of heat-shock genes (2), an apparently unique feature that is not universal across the genus *Drosophila* (3).

The artificial P elements have been broadly used for genetic analysis and functional genomics in *D. melanogaster* and are a valuable resource for carrying out targeted and random mutagenesis (4). Concerning *P{lacW}* and *P{EP}* artificial TEs, they display a series of structural and functional similarities and differences (5, 6). The length of the *P{lacW}* mobile element is 10,691 bp and it harbors at the 5’ end the *lacZ* coding sequences, inside the central region it carries the *Hsp70* promoter and the *mini-white* gene, while *ampR* is sited at its 3’ end (5, 7). The *P{EP}* mobile element has a length of 7,987 bp and consists of *mini-white*, located at the 5’ end, a central *kanR* and within the 3’ end the UAS regulatory sequences, *Hsp70* and GAL4-inducible promoters (6, 7). Since both *P{lacW}* and *P{EP}* derive from the natural P element, they have the same 31 bp terminal inverted repeats and neighboring internal sequences, which are required for transposase binding and transposition.

It is of interest to investigate the degree to which the insertion patterns of these related TEs are affected because of their size and cargo differences, especially considering that they are sharing the same transposition mechanisms.

In this study, we analyzed the spatial and numerical distribution of *P{lacW}* and *P{EP}* TEs within the genome of *D. melanogaster*, exploiting the insertional datasets available in FlyBase.

## Materials and methods

The *insertion_mapping_fb_2019_04.tsv.gz* file, which contains data regarding the insertion occurrences of various types of artificial TEs, was downloaded from the FlyBase database (http://flybase.org). The file is available at *Downloads* section, the *Transposons, Transgenic Constructs, and Insertions* category. The tabular data from the *insertion_mapping_fb_2019_04.tsv.gz* file contain the symbols of TEs and, when available, the genomic location, the orientation of TEs, as well as the estimated and/or observed cytogenetic location.

We used *grep* command to extract from the unzipped file data referring only to *P{lacW}* and *P{EP}*.

*$ grep “P{lacW}” insertion_mapping_fb_2019_04.tsv* > *PlacW_insertions.tsv*

*$ grep “P{EP}” insertion_mapping_fb_2019_04.tsv* > *PEP_insertions.tsv*

The resulting files, symbolized *PlacW_insertions.tsv* and *PEP_insertions.tsv*, contain several hundreds of TE insertions without any genomic coordinate or cytogenetic location, which were not considered for further analyses. Data filtering was performed with *LibreOffice Calc* version 6.0.7.3 and *Bash* command line on Ubuntu 18.04.2 LTS. The filtered *PlacW_insertions.tsv* and *PEP_insertions.tsv* files contain 5,581 *P{lacW}* and 3,787 *P{EP}* insertions. Since a number of 434 *P{lacW}* and 558 *P{EP}* insertions did not have a standard representation of the estimated cytogenetic location. In order to use them for the purpose of this study, we selected and loaded the FlyBase IDs of these insertions to the *Batch Download* tool from FlyBase and consequently obtained standard estimated cytogenetic locations. A number of 19 TEs insertions (one *P{EP}* and 18 *P{lacW}*s) were not associated with a unique chromosomal division (region), therefore they were sorted out from the final data.

The remaining TEs insertions were assigned to corresponding chromosomal divisions based on their estimated cytogenetic locations by using *table* function in RStudio. Therefore, we obtained *Frequency tables* files that contain the number of occurrences of every insertion in each division of the X chromosome and 2L, 2R, 3L, and 3R chromosome arms.

We did not map any insertion on chromosome Y, while for chromosome 4 we only found three *P{lacW}* insertions within 101F, 102A1 and 102D5-102D6 sections. These data were not further analyzed.

The plots and statistical analysis (nonparametric Mann-Whitney test and Spearman correlation) were performed using GraphPad Prism 5.04 software (GraphPad Software, La Jolla California USA, www.graphpad.com).

Fourier analysis was performed with the R programming language. Fast Fourier Transform was performed using the function “*fft*” and the interpolation with “*pracma*” package’s “*interp1*”.

## Results

A total number of 5,560 *P{lacW}* and 3,786 *P{EP}* insertions were asigned to their corresponding chromosomes (Table 1). More than half of the total number of *P{lacW}* insertions (52.93%) are located on the chromosome 2, while the majority of *P{EP}* insertions (42.92%) are placed on chromosome 3.

**Table 1.**
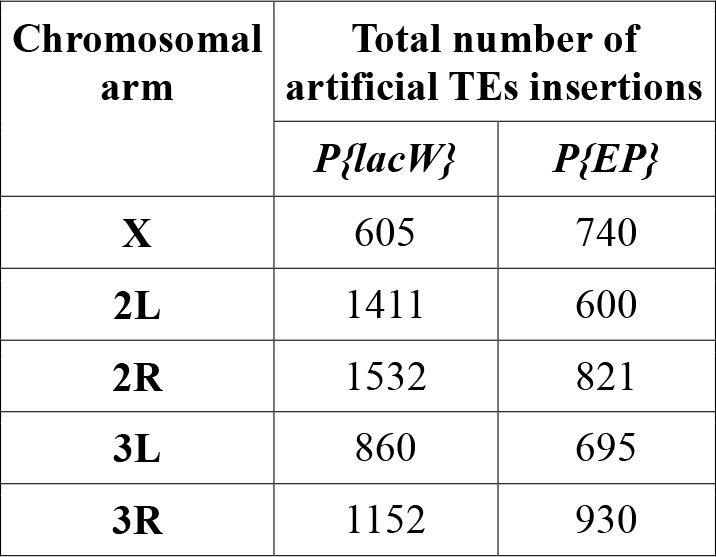
The distribution of *P{lacW}* and *P{EP}* insertions inside the X chromosome and 2L, 2R, 3L, and 3R chromosome arms.

| Chromosomal arm | Total number of artificial TEs insertions |  |
| --- | --- | --- |
|  | *P{lacW}* | *P{EP}* |
| **X** | 605 | 740 |
| **2L** | 1411 | 600 |
| **2R** | 1532 | 821 |
| **3L** | 860 | 695 |
| **3R** | 1152 | 930 |

To further refine our analysis, we allocated the *P{lacW}* and *P{EP}* insertions to chromosomal divisions and consecutively plotted the spatial and numeric distribution patterns of both TEs on the corresponding chromosome. For each chromosome we pinpointed both the division harboring a maximum of *P{lacW}* or *P{EP}* insertions and the top three hotspot or refractory divisions for insertional events, distinctly for *P{lacW}* and *P{EP}*.

On chromosome X, we noticed that there are both similarities and differences between *P{lacW}* and *P{EP}* distribution (Figure 1). For both TEs, the lowest number of insertions is scored on the last divisions of the chromosome (division 20), with only three insertions of *P{lacW}* and four of *P{EP}*. Division 12 has the highest number of *P{EP}* insertions (93 insertions), while *P{lacW}* insertions are enriched for division 3 (67 insertions).

**Figure 1.**
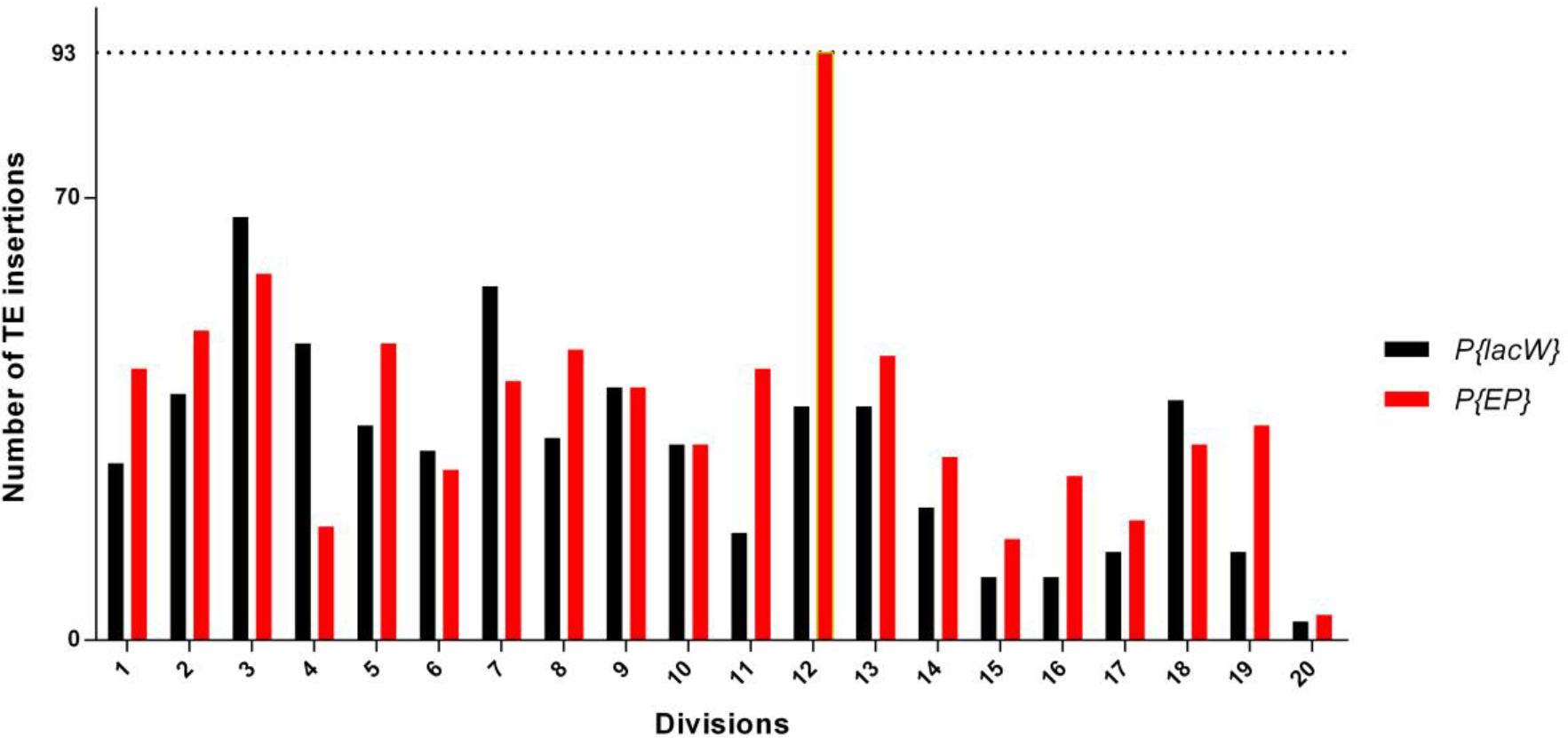
Comparison of insertion distribution of *P{lacW}* (black bars) and *P{EP}* (red bars) on the X chromosome of *D. melanogaster*. The x-axis shows the divisions of the X chromosome and on the y-axis are represented the number of TE insertions. With yellow border is highlighted the maximal insertional load that is reached by *P{EP}* on division 12 (93 insertions).

When considering *P{lacW}*, the top three hotspot and refractory regions are 3, 4, and 7, while the most refractory regions for insertional events are 15, 16, and 20. Regarding *P{EP}*, the hotspot divisions are 2, 3 and 12, and refractory divisions are 15, 17 and 20 (Table 2). It worth mentioning that the hotspot division 3 and refractory divisions 15 and 20 are common to both *P{lacW}* and *P{EP}*.

**Table 2.**
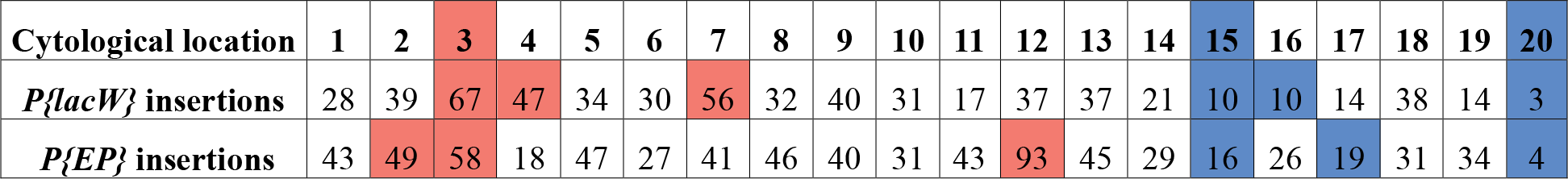
Number of *P{lacW}* and *P{EP}* insertions/division of X chromosome. We highlighted in red the top three hotspot divisions and in blue the top three refractory divisions.

| Cytological location | 1 | 2 | 3 | 4 | 5 | 6 | 7 | 8 | 9 | 10 | 11 | 12 | 13 | 14 | 15 | 16 | 17 | 18 | 19 | 20 |
| --- | --- | --- | --- | --- | --- | --- | --- | --- | --- | --- | --- | --- | --- | --- | --- | --- | --- | --- | --- | --- |
| $P\{lacW\}$ insertions | 28 | 39 | 67 | 47 | 34 | 30 | 56 | 32 | 40 | 31 | 17 | 37 | 37 | 21 | 10 | 10 | 14 | 38 | 14 | 3 |
| $P\{EP\}$ insertions | 43 | 49 | 58 | 18 | 47 | 27 | 41 | 46 | 40 | 31 | 43 | 93 | 45 | 29 | 16 | 26 | 19 | 31 | 34 | 4 |

The chromosome arm 2L displays a different distribution for *P{lacW}* and *P{EP}* as compared to chromosome X (Figure 2). Similar to X, the lowest number of insertions scored for both TEs are on the last division of the chromosome arm (division 40), adjacent to centromeric region, with 25 insertions of *P{lacW}* and three of *P{EP}*. The maximum number of insertions is scored for *P{lacW}* in division 35 (152 insertions). The 21 division, including the telomeric end, represents a common hotspot for both TEs, while divisions 24 and 40 are common refractory regions.

**Figure 2.**
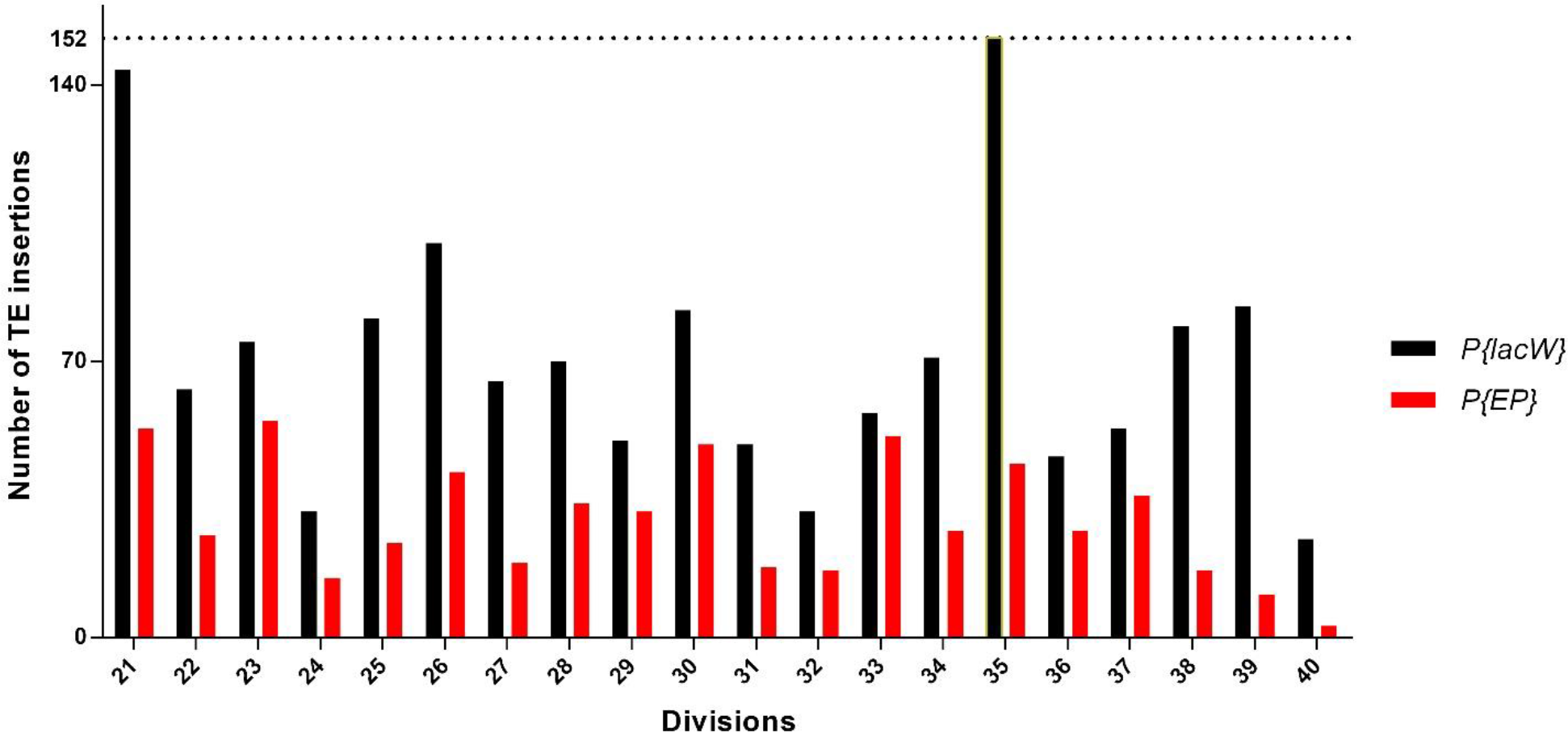
Comparison of insertion distribution of *P{lacW}* (black bars) and *P{EP}* (red bars) on chromosome arm 2L of *D. melanogaster*. The x-axis shows the divisions of the 2L chromosome arm and on the y-axis are represented the number of TE insertions. With yellow border is highlighted the maximal insertional load that is reached by *P{lacW}* on division 35 (152 insertions).

For *P{lacW}*, the top three hotspot and refractory regions are 21, 26, and 35, and, respectively, 24, 32, and 40, while regarding *P{EP}*, hotspot divisions are 21, 23 and 33, and refractory divisions are 24, 39 and 40 (Table 3).

**Table 3.** Number of *P{lacW}* and *P{EP}* insertions/division of chromosome arm 2L. We highlighted in red the top three hotspot divisions and in blue the top three refractory divisions.

| Cytological location | 21 | 22 | 23 | 24 | 25 | 26 | 27 | 28 | 29 | 30 | 31 | 32 | 33 | 34 | 35 | 36 | 37 | 38 | 39 | 40 |
| --- | --- | --- | --- | --- | --- | --- | --- | --- | --- | --- | --- | --- | --- | --- | --- | --- | --- | --- | --- | --- |
| $P\{lacW\}$ insertions | 144 | 63 | 75 | 32 | 81 | 100 | 65 | 70 | 50 | 83 | 49 | 32 | 57 | 71 | 152 | 46 | 53 | 79 | 84 | 25 |
| $P\{EP\}$ insertions | 53 | 26 | 55 | 15 | 24 | 42 | 19 | 34 | 32 | 49 | 18 | 17 | 51 | 27 | 44 | 27 | 36 | 17 | 11 | 3 |

The *P{lacW}* and *P{EP}* distribution on chromosome arm 2R exhibits similarities with 2L (Figure 3). Thus, for both TEs, the lowest number of insertions was counted for division 41, which is next to the centromeric region and harbors 12 insertions of *P{lacW}* and four of *P{EP}*. Interestingly, the highest number of insertions is totaled for *P{lacW}* within division 42 (118 insertions), positioned relatively close to the centromeric region. At the opposite end, the divisions 57 and 60 are hotspots for both TEs, and division 59 is a refractory region for *P{lacW}* and *P{EP}* (Table 4).

**Table 4.** Number of *P{lacW}* and *P{EP}* insertions/division of chromosome arm 2R. We highlighted in red the top three hotspot divisions and in blue the top three refractory divisions.

| Cytological location | 41 | 42 | 43 | 44 | 45 | 46 | 47 | 48 | 49 | 50 | 51 | 52 | 53 | 54 | 55 | 56 | 57 | 58 | 59 | 60 |
| --- | --- | --- | --- | --- | --- | --- | --- | --- | --- | --- | --- | --- | --- | --- | --- | --- | --- | --- | --- | --- |
| $P\{lacW\}$ insertions | 12 | 118 | 56 | 80 | 76 | 79 | 108 | 76 | 86 | 101 | 63 | 59 | 90 | 60 | 81 | 68 | 116 | 40 | 46 | 117 |
| $P\{EP\}$ insertions | 4 | 35 | 39 | 24 | 42 | 34 | 62 | 33 | 48 | 39 | 31 | 34 | 55 | 51 | 45 | 27 | 61 | 28 | 27 | 102 |

**Figure 3.**
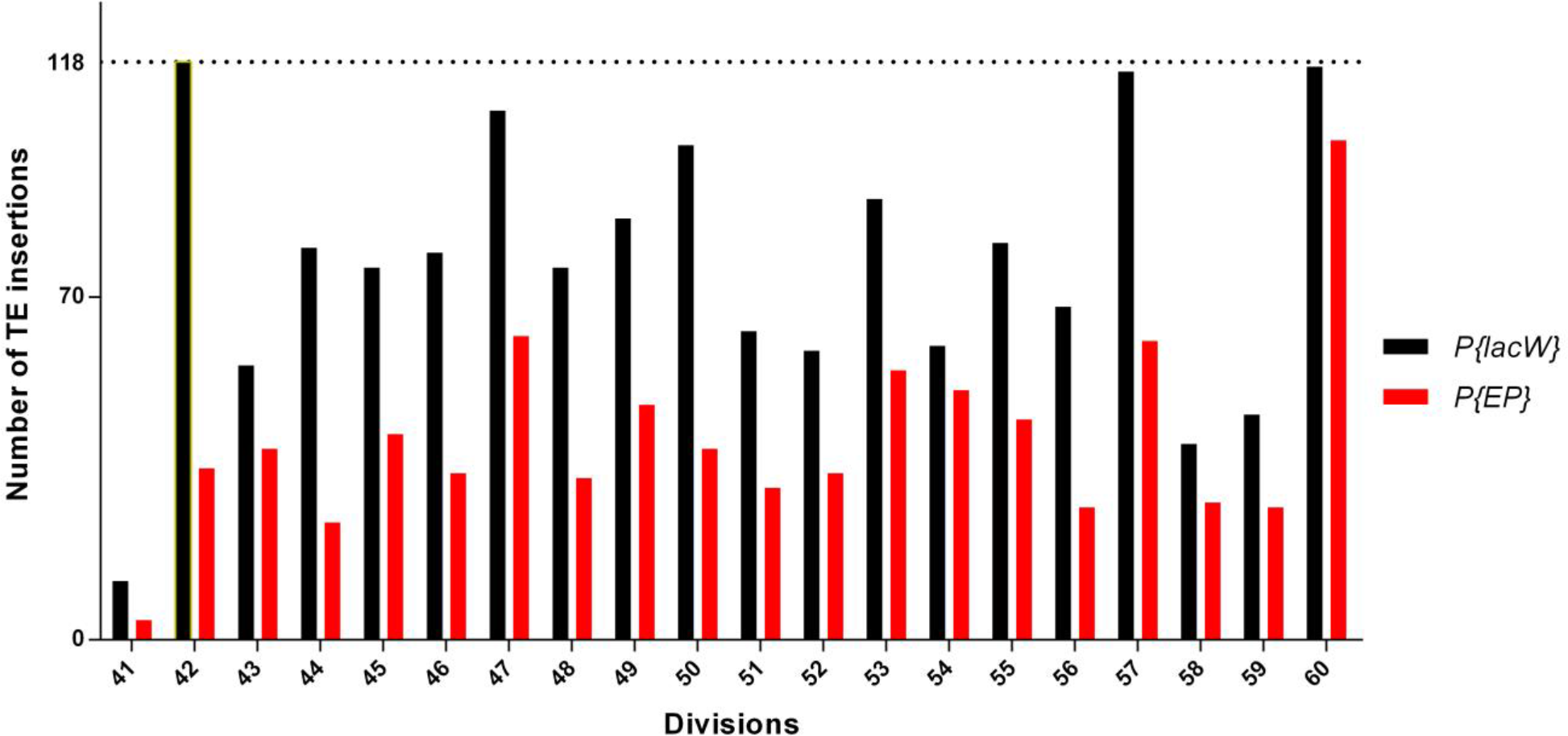
Comparison of insertion distribution of *P{lacW}* (black bars) and *P{EP}* (red bars) on chromosome arm 2R of *D. melanogaster*. The x-axis shows the divisions of the 2R chromosome arm and on the y-axis are represented the number of TE insertions. With yellow border is highlighted the maximal insertional load that is reached by *P{lacW}* on division 42 (118 insertions).

The top three hotspot and refractory regions for *P{lacW}* are 42, 57, and 60, and, respectively, 41, 58, and 59. For *P{EP}*, hotspot divisions are 47, 57 and 60, and refractory divisions are 41, 44, 56, and 59, the last two divisions having an equal number of insertions (Table 4).

Aside from chromosome X, the chromosome arm 3L harbors the lowest total number of *P{lacW}* and *P{EP}* insertions (Figure 4). Division 75 is highly enriched for *P{lacW}* insertions (84 insertions), but division 66 (common hotspot) accumulates a maximum number of *P{lacW}* and *P{EP}* insertions (a total of 141 insertions). The lowest number of insertions is recorded in division 80, which contains eight *P{lacW}* and six *P{EP}* insertions. Another common refractory is division 74, with a total of 27 insertions.

**Figure 4.**
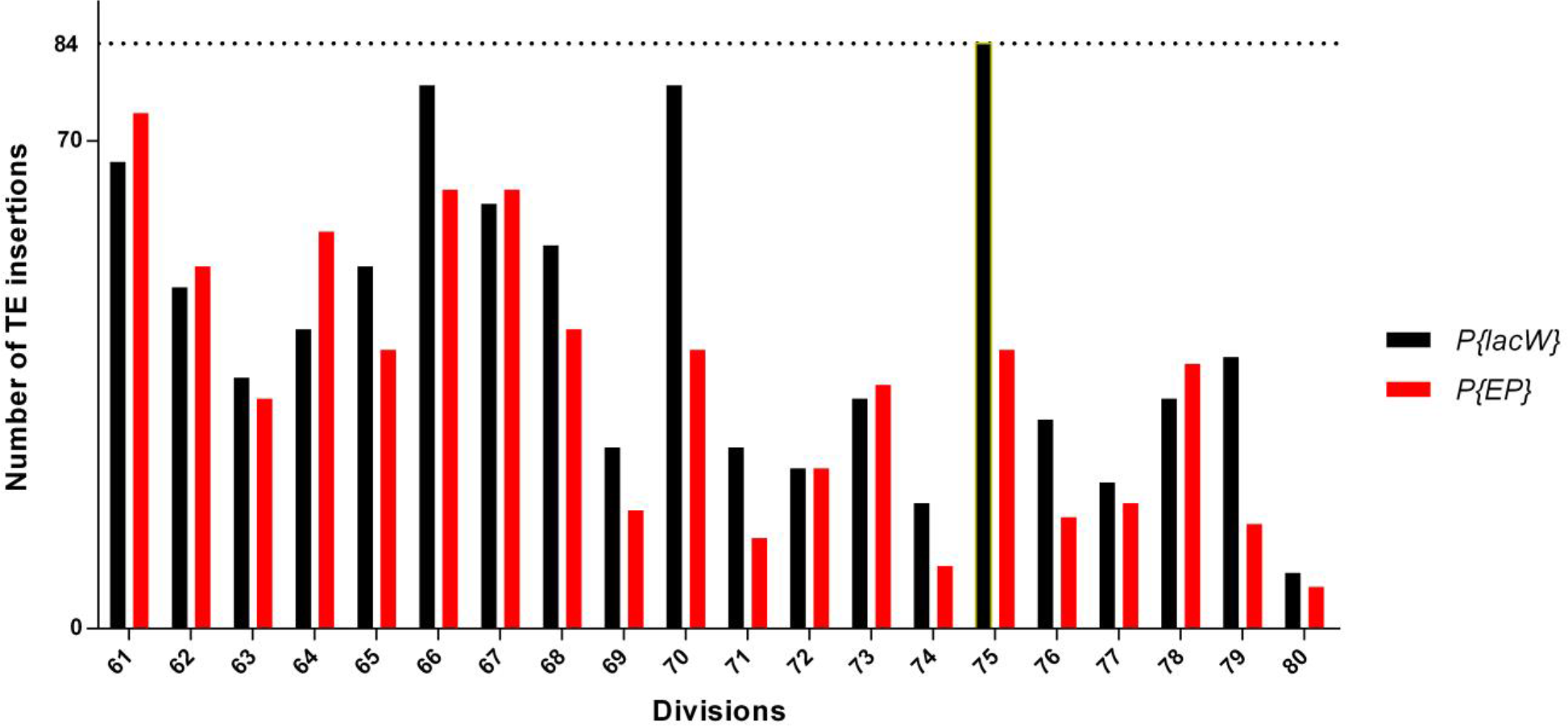
Comparison of insertion distribution of *P{lacW}* (black bars) and *P{EP}* (red bars) on 3L chromosome arm of *D. melanogaster*. The x-axis shows the divisions of the 3L chromosome arm and on the y-axis are represented the number of TE insertions. With yellow border is highlighted the maximal insertional load that is reached by *P{lacW}* on division 75 (84 insertions).

The top three hotspot and refractory regions for *P{lacW}* are 66, 70, and 75, and, respectively, 74, 77, and 80. For *P{EP}*, hotspot divisions are 61 (telomeric end), 66 and 67, and refractory divisions are 71, 74, and 80 (Table 5).

**Table 5.** Number of *P{lacW}* and *P{EP}* insertions/division of chromosome arm 3L. We highlighted in red the top three hotspot divisions and in blue the top three refractory divisions.

| Cytological location | 61 | 62 | 63 | 64 | 65 | 66 | 67 | 68 | 69 | 70 | 71 | 72 | 73 | 74 | 75 | 76 | 77 | 78 | 79 | 80 |
| --- | --- | --- | --- | --- | --- | --- | --- | --- | --- | --- | --- | --- | --- | --- | --- | --- | --- | --- | --- | --- |
| $P\{lacW\}$ insertions | 67 | 49 | 36 | 43 | 52 | 78 | 61 | 55 | 26 | 78 | 26 | 23 | 33 | 18 | 84 | 30 | 21 | 33 | 39 | 8 |
| $P\{EP\}$ insertions | 74 | 52 | 33 | 57 | 40 | 63 | 63 | 43 | 17 | 40 | 13 | 23 | 35 | 9 | 40 | 16 | 18 | 38 | 15 | 6 |

Comparative to other chromosome arms, where the frequency of *P{lacW}* insertions are evidently greater than those particular for *P{EP}*, 3R displays a more balanced proportion between the two types of insertions (Figure 5). Regarding both *P{lacW}* and *P{EP}* insertions, 3R contains two common refractory divisions (the peri-centromeric division 81, with only one *P{EP}* insertion, and division 97). The highest number of insertions are counted in division 85 for both *P{lacW}* and *P{EP}* insertions (127 and, respectively, 106 insertions).

**Figure 5.**
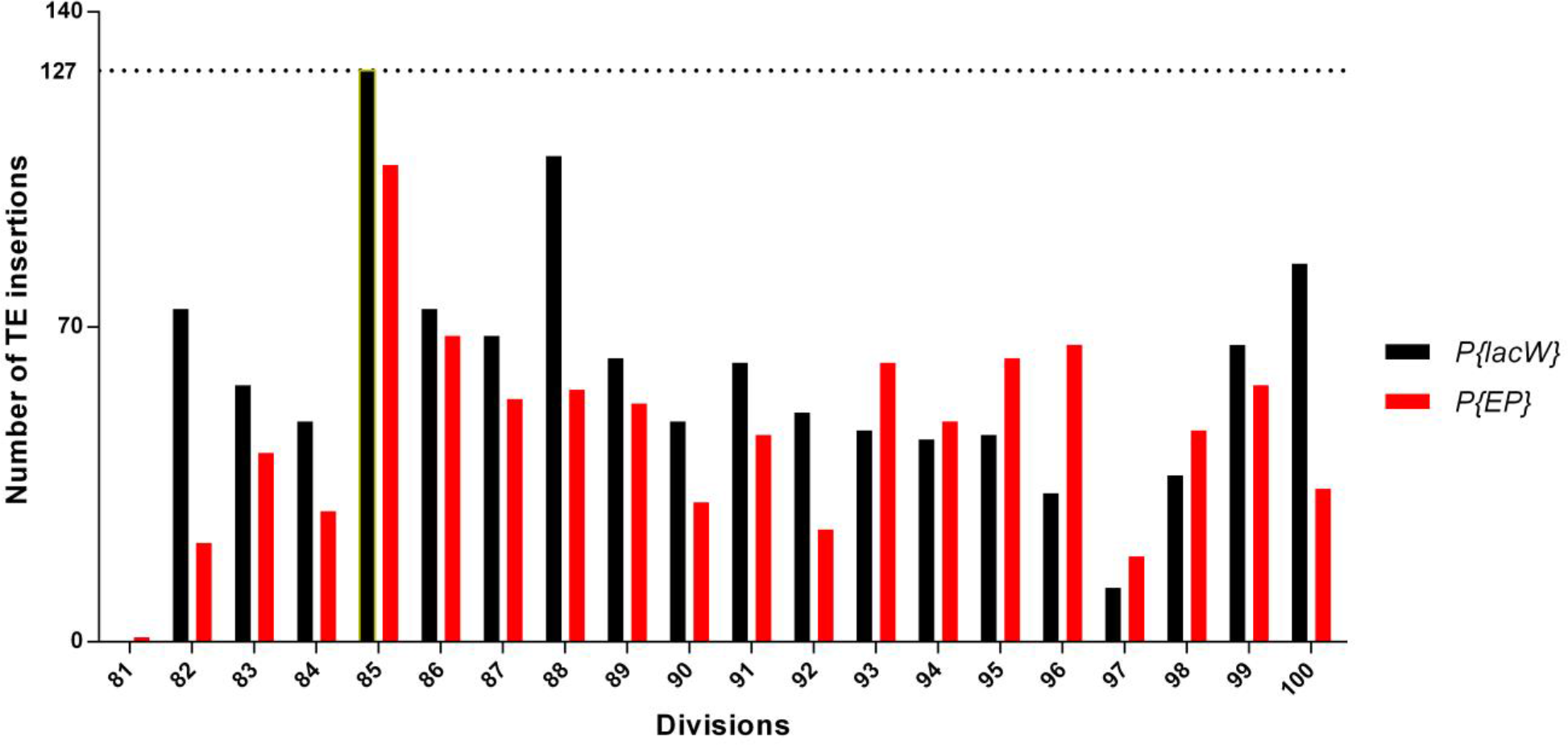
Comparison of insertion distribution of *P{lacW}* (black bars) and *P{EP}* (red bars) on 3R chromosome arm of *D. melanogaster*. The x-axis shows the divisions of the 3R chromosome arm and on the y-axis are represented the number of TE insertions. With yellow border is highlighted the maximal insertional load that is reached by *P{lacW}* on division 85 (127 insertions).

The three most distinctive hotspot and refractory regions for *P{lacW}* are 85, 88, and 100 (telomeric end), and, respectively, 81, 96, and 97. For *P{EP}*, hotspot divisions are 85, 86 and 96, and refractory divisions are 81, 82, and 97 (Table 6).

**Table 6.** Number of *P{lacW}* and *P{EP}* insertions/division of chromosome arm 3R. We highlighted in red the top three hotspot divisions and in blue the top three refractory divisions.

| Cytological location | 81 | 82 | 83 | 84 | 85 | 86 | 87 | 88 | 89 | 90 | 91 | 92 | 93 | 94 | 95 | 96 | 97 | 98 | 99 | 100 |
| --- | --- | --- | --- | --- | --- | --- | --- | --- | --- | --- | --- | --- | --- | --- | --- | --- | --- | --- | --- | --- |
| $P\{lacW\}$ insertions | 0 | 74 | 57 | 49 | 127 | 74 | 68 | 108 | 63 | 49 | 62 | 51 | 47 | 45 | 46 | 33 | 12 | 37 | 66 | 84 |
| $P\{EP\}$ insertions | 1 | 22 | 42 | 29 | 106 | 68 | 54 | 56 | 53 | 31 | 46 | 25 | 62 | 49 | 63 | 66 | 19 | 47 | 57 | 34 |

The insertion distributions graphics, as well as the corresponding tabular data, indicates that *P{lacW}* and *P{EP}* may have a similar insertional behavior (our null hypothesis *H*_*0*_ states that there is a significant difference between the two patterns of insertion). In order to test *H*_*0*_, we transformed the numerical values representative for each type of insertion/chromosomal division in fraction of the total insertions/chromosome arm. The values corresponding to *P{lacW}* and those corresponding to *P{EP}* insertions located on a certain chromosome arm were compared using *Mann-Whitney* test. All comparisons provided P values that are not significant, therefore there are no differences between the fractions of these TEs insertions within the divisions of a certain chromosome (for X, P = 0.9627; for 2L, P > 0.9999; for 2R, P = 0.5969; for 3L, P = 0.8567, for 3R, P = 0.9627). In addition, we evaluated the similarities between the distributions of *P{lacW}* and *P{EP}* insertions by applying Spearman correlation. We found that there is a significant correlation between the distribution of *P{lacW}* and *P{EP}* for chromosomes X, and whole chromosome 2 and chromosome 3 (Figure 6). All correlations are positive and statistically significant with the following values: for chromosome X, r = 0.5582 and P = 0.0105; for chromosome 2, r = 0.5966 and P < 0.0001; for chromosome 3, r = 0.6235 and P < 0.0001. When we calculated Pearson’s correlation coefficient for chromosomal arms, data were significant and positive correlated for 2L (r = 0.4829 and P = 0.0310), 2R (r = 0.6574 and P = 0.0016), 3L (r = 0.8262 and P < 0.0001), but not for 3R (r = 0.3138 and P = 0.1779).

**Figure 6.**
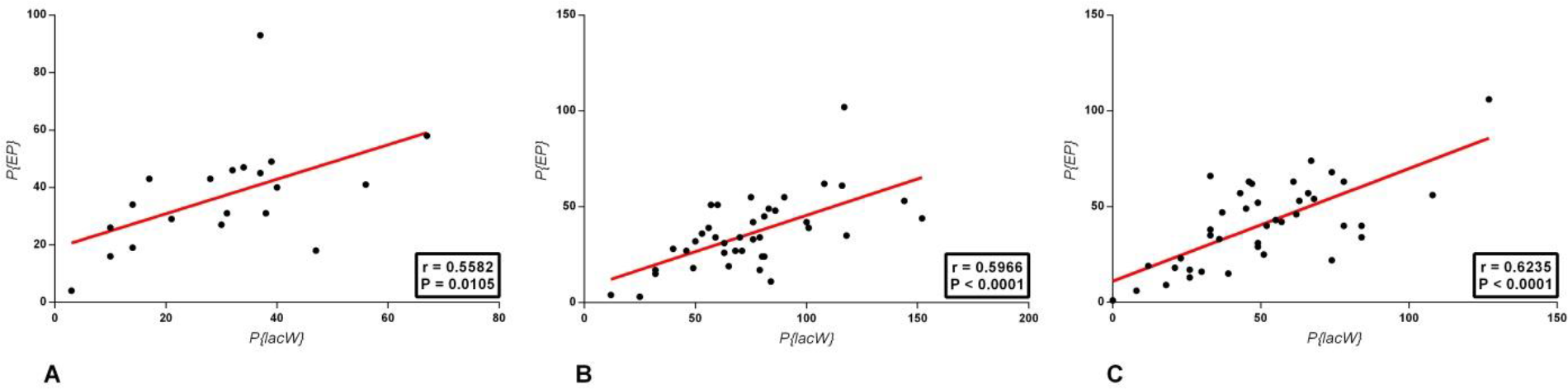
Spearman correlations for chromosome X (A), chromosome 2 (B) and chromosome 3 (C). For each chromosome, the distributions of *P{lacW}* and *P{EP}* insertions, the linear regression (in red) and a panel shoving the related r and P values are represented.

To reveal any periodic behaviour in the distributions of insertions, we used the Fast Fourier Transform (8) to obtain their spectra. For a proper usage of the algorithm, the number of data points had to be artificially increased from 20 (the number of divisions/chromosome arm) to 1000 using spline interpolation (9). Frequency peaks (a peak is a local maxima where the value is greater than the values on the left and on the right of the plot) were marked and the peaks which were common to both *P{lacW}* and *P{EP}* insertions on the same chromosome were highlighted (Figure 7). Since the distribution of *P{lacW}* and *P{EP}* insertions on the same chromosomes are significantly correlated (excepting for 3R), the common frequency peaks suggest a periodical occurrence of insertional hotspots. A reliable periodic distribution is expected to have only a few outstanding peak frequencies as can be seen in X for *P{lacW}*, while a rather noisy periodic distribution has many weak peaks, as for *P{EP}* in 2R. On X chromosome, *P{EP}* insertions are noisy, since the spectrum is made of many frequencies with similar amplitudes. Nevertheless, the value 4 specific for *P{lacW}* insertions frequency on the same chromosome is still present among them. This aspect suggests that *P{EP}* distribution on X chromosome is a sum of a harmonic with frequency common with *{PlacW}* (the highlighted one in Figure 7, X P{lacW} and X P{EP} frames) and of noise.

**Figure 7.**
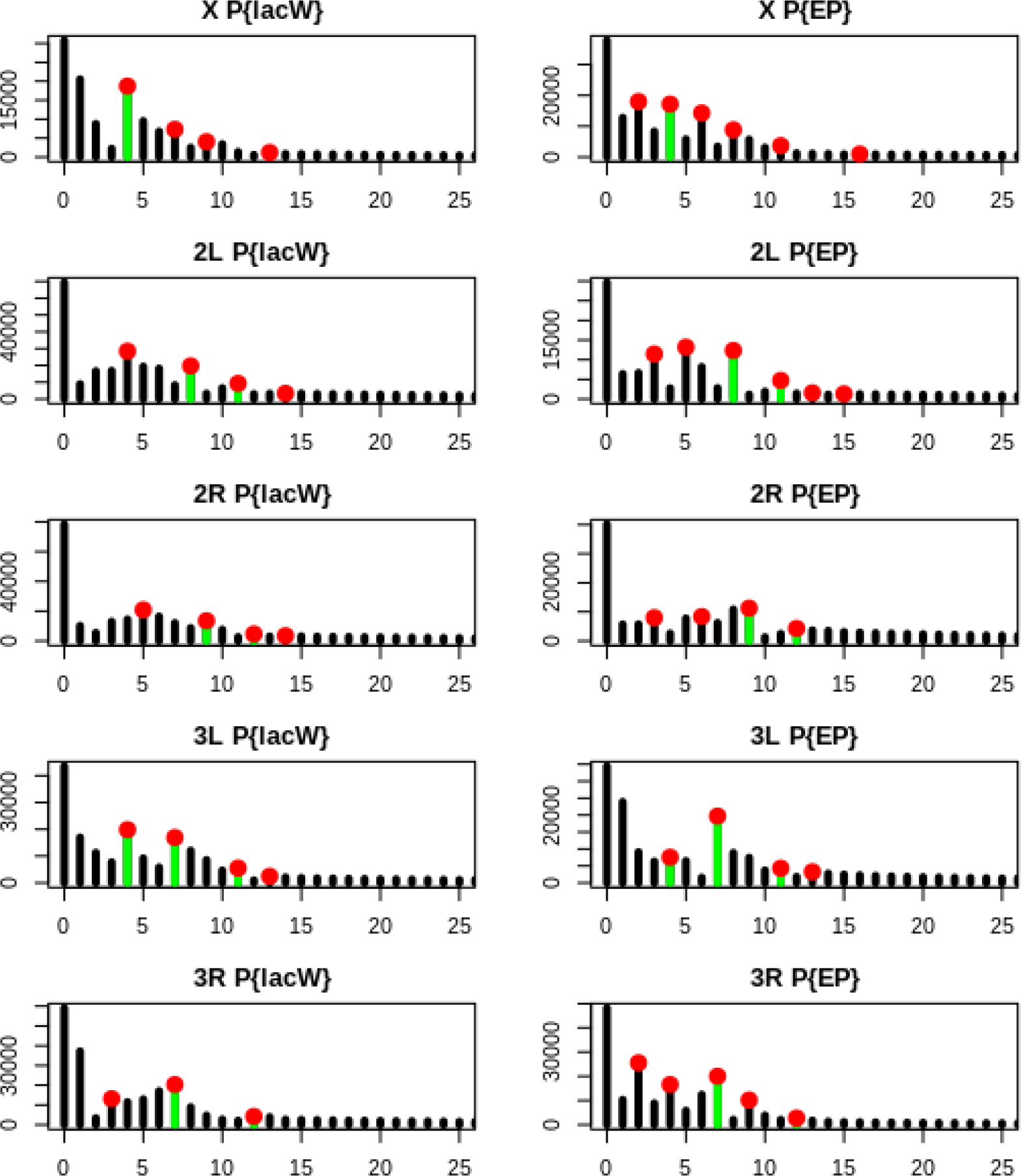
Spectra for the regional insertion patterns of *P{lacW}* and *P{EP}* on each chromosome arm. The *x* axis is the wavelength, the *y* axis is the relative intensity of a particular wavelength (the contribution of the periodic with that wavelength to the overall distribution). Peak wavelengths which are local maxima are marked with a red dot. Peak wavelengths common to both TEs on the same chromosome arm are highlighted in green.

The strongest common periodic behavior is in 3L, where both spectra are dominated by two peak frequencies. 2R is both the weakest periodic and has the weakest common frequencies between the two transposons. In each chromosome, one common peak between the transposons can be identified which could be suggestive for a particular insertion pattern on that chromosome. The wavelength, in this case, is defined as 20 ÷ *frequency*.

In X, the common wavelength is 4, with the first hotspot in chromosomal division 3, thus the next hotspots should be around the regions 8, 13 and 18 (see Figure 1). Similarly, the common wavelength in 2L is 2.5, in 2R there is a weak wavelength of 2.22, and in 3L and 3R it is 2.86.

## Discussions

The distribution of *P{lacW}* insertions obtained in our study for chromosome 3 of *D. melanogaster* is consistent with the data from previous studies (Deak *et al.*, 1997; Popa *et al.*, 2016). To our best knowledge, we are not aware of other similar studies focusing on a comparative analysis of the distribution of related artificial TEs insertions. Our results generated by *Mann-Whitney* test and *Spearman* correlation analysis suggest that the insertion’s distribution of *P{lacW}* and *P{EP}* artificial TEs are similar for the chromosomes X, 2 and 3 of *D. melanogaster*. This aspect may be explained by the fact that both *P{lacW}* and *P{EP}* are derived from the natural P element, therefore their transposition mechanism is expected to be the same. Obviously, the two distributions of insertions are not identical, probably because of their diferent molecular lenght and of specific transgenes cargo. Other factors, such as the specific conditions of the insertional mutagenesis experiments, may also be responsible for the observed differences of distributions. *Fourier* analysis suggests a periodic behaviour of the two TEs insertion’s distribution, which approximates the insertional hotspots.

Since the *P{lacW}* and *P{EP}* insertion patterns are similar to each other, it is tempting to assume that each pattern is not random *per se*, reflecting insertional preferences. This aspect is in agrement with other data concerning insertional preferences of natural transposons (Bergman ….).

## Conclusions

Our preliminary results raise questions concerning if and how the local chromosomal landscape affects the insertion patterns of different but related transposons. It is also tempting to find out if the insertions of artificial and natural transposons of class II in the genome of *D. melanogaster* display similarities of insertional patterns.

